# Two new genetically modified mouse alleles labeling distinct phases of retinal ganglion cell development by fluorescent proteins

**DOI:** 10.1101/2020.03.03.975144

**Authors:** Yichen Ge, Fuguo Wu, Mobin Cheng, Xiuqian Mu

## Abstract

During development, retinal progenitor cells (RPCs) take on different trajectories for distinct cell fates. Previous studies have identified key regulators involved in two key steps of retinal ganglion cell (RGC) genesis; Atoh7 functions in a subpopulation of RPCs to render them competent for the RGC fate, whereas Pou4f2 and Isl1 function to specify the RGC fate and promote RGC differentiation. Extensive research has been performed on the roles of these transcription factors in RGC development, but properties of these two phases they represent and the cellular context in which these two factors function have not been thoroughly investigated, largely due to the cellular heterogeneity of developing retina. In this paper, we describe two novel knock-in mouse alleles, *Atoh7*^*zsGreenCreERT2*^ and *Pou4f2*^*FlagtdTomato*^, which enabled us to label retinal cells in the two phases of RGC development by fluorescent proteins. In addition, the *Atoh7*^*zsGreenCreERT2*^ allele also allowed for indirect labeling of RGCs and other cell types upon tamoxifen induction in a dose-dependent manner. Further, these alleles could be used to purify retinal cells of these different phases by fluorescence assisted cell sorting (FACS). Thus, these two alleles can serve as very useful tools for studying the molecular and genetic mechanisms underlying RGC formation.

## Introduction

The neural retina serves as a good model for studying the genetic and molecular mechanisms governing the formation of cellular diversity in the central nervous system. There are seven major cell types in the neural retina, all derived from a single pool of retinal progenitor cells (RPCs) during development (Cepko et al., 1996). Generation of these cell types follows a distinct temporal order in all vertebrate species due to changes in competence of RPCs over time (Cepko, 2014; Cepko et al., 1996; Young, 1985). On the other hand, formation of each cell type is governed by a specific gene regulatory cascade in a step-wise fashion (Bassett and Wallace, 2012; Swaroop et al., 2010; Xiang, 2013). Much has been learned about the key regulators (transcription factors) and their roles in the generation of individual cell types. However, these studies have largely been performed at the whole-retina level, and thus, due to the heterogeneity of the developing retina, the properties of cells in the different phases of the developmental trajectory of individual cell lineages and the cellular context in which the transcription factors function have not been well addressed.

Retinal ganglion cells (RGCs) are the only projection neuron in the retina, which emit axons in the form of the optic nerve to the targets in the brain (Dhande and Huberman, 2014; Masland, 2001; Sanes and Masland, 2015; Seabrook et al., 2017). RGCs are the first cell type to be born, although its generation has significant overlap with the other early cell types, including horizontal cells, amacrine cells, and cones (Cepko et al., 1996; Young, 1985). In the mouse retina, RGC formation starts at embryonic day (E) 11.5 and reaches the peak at E14.5 (Young, 1985). Like other cell types, RGC genesis is a step-wise process. Key transcription factors essential for RGC development have been identified, and their functions have been studied. Among them are Atoh7, which functions in a subset of RPCs, and Pou4f2 and Isl1, which function in RGCs and their precursors (Brown et al., 2001; Gan et al., 1996; Mu et al., 2008; Pan et al., 2008; Wang et al., 2001). Consistent with their functions in RGCs, expression of these factors largely follows the same temporal dynamics of RGC production (Brown et al., 1998; Fu et al., 2009; Miesfeld et al., 2018; Xiang et al., 1995). Atoh7 is essential, but not sufficient, for the RGC lineage; deletion of *Atoh7* leads to failed RGC production, but *Atoh7*-expressing cells adopt multiple cell fates (Brown et al., 2001; Brzezinski et al., 2012; Feng et al., 2010; Wang et al., 2001; Yang et al., 2003). Deletion of *Pou4f2* or *Isl1* leads to aberrant RGC differentiation, and ectopic expression of Pou4f2 and Isl1 can rescue RGC production in the *Atoh7*-null background (Mu et al., 2008; Pan et al., 2008; Wu et al., 2015). Thus, Atoh7 renders RPCs competent for the RGC fate, whereas Pou4f2 and Isl1 specify the RGC fate and promote their further differentiation.

Transcriptomic analyses have provided much insight into how these factors exert their functions and what target genes they regulate (Mu et al., 2004, 2005, 2008; Qiu et al., 2008). However, since these studies have been carried out at the whole-retina level, it has not been possible to investigate the molecular properties of the cell states in which these transcription factors operate, including genes being expressed and the underlying epigenetic landscape responsible for their expression. For example, it is not known what makes the *Atoh7*-expressing RPC competent for the RGC fate and what is their relationships to RPCs relevant to other cell fates. It is also not known what happens to the epigenetic landscape during the transition from multipotent RPCs to fate-committed RGCs. These questions may only be addressed with purified cells representing the different phases of RGC development. With the advantages of being highly sensitive and non-destructive to the function of target cells and proteins, fluorescent protein tags are widely used in labeling and tracking cells. Genetic labeling by fluorescent proteins also enables isolation and study of particular cell populations during development. In this report, we have generated two fluorescent protein-tagged knock-in mouse alleles, *Atoh7*^*zsGreenCreERT2*^ and *Pou4f2*^*FlagtdTomato*^. These alleles labeled retinal cells in the two critical phases of RGC development respectively by two fluorescent proteins, zsGreen and tdTomato. The *Atoh7*^*zsGreenCreERT2*^ also could be used to label RGCs indirectly with a conditional reporter allele and to delete other floxed alleles for functional analysis in a dose-dependent fashion upon induction by tamoxifen. We envision that these two alleles will serve as very useful tools for studying the molecular and genetic mechanisms underlying RGC formation.

## Materials and methods

### Animals

The Cre-dependent tdTomato reporter mouse Ai9 (B6.Cg-*Gt (ROSA)26Sor*^*tm9(CAG-tdTomato)Hze*^/J, 007909) was obtained from the Jackson Laboratory (Madisen et al., 2010). The *Atoh7*^*HA*^ and *Atoh7*^*laz*^ alleles have been described previously (Fu et al., 2009; Wang et al., 2001). All mice in the current study were maintained in a C57/BL6×129 genetic background. All animal procedures conformed to the US Public Health Service Policy on Humane Care and Use of Laboratory Animals and were approved by the Institutional Animal Care and Use Committees of Roswell Park Comprehensive Cancer Center and the University at Buffalo.

### Gene Targeting

Gene targeting constructs for *Atoh7* ^*zsGreenCreERT2*^ and *Pou4f2*^*FlagtdTomato*^ were generated by recombineering following the procedure of Liu et al., 2003 (Liu et al., 2003). The general strategy to generate the targeting constructs were the same as previously reported for the HA-tagged alleles of the two genes (Fu et al., 2009). To generate the *Atoh7*^*zsGreenCreERT2*^ targeting vector, a cassette containing the following components, *zsGreen*–*T2A*–*CreERT2*–SV40 poly(A)–*LoxP*–*Neo*–*LoxP*, was constructed by Gibson cloning (Gibson et al., 2009). This cassette was then used to replace the *Atoh7* ORF in a genomic DNA fragment captured by BAC recombineering (Fu et al., 2009). The final construct contains a 5.3 kb 5’ arm, the *zsGreen*–*T2A*–*CreERT2*–SV40 poly(A)–*LoxP*– *Neo*–*LoxP* cassette (4.8 kb), and a 1.8 kb 3’ arm, and a TK cassette, and could be linearized by a single NotI site.

A similar procedure was used to construct the *Pou4f2*^*flagtdTomato*^ targeting vector. A cassette containing the following components, *FLAG*–*T2A*–*tdTomato*–SV40 poly(A)– *LoxP*–*Neo*–*LoxP*, was generated by Gibson cloning. The cassette was then cloned in frame after the last codon of Pou4f2 in the genomic DNA fragment captured previously (Fu et al., 2009). This targeting construct contained a 4.3 kb 5’ arm with the two exons encoding the full-length Pou4f2, the *FLAG*–*T2A*–*tdTomato*–SV40 poly(A)–*LoxP*–*Neo*– *LoxP* cassette (3.6 kb), a 3.1 kb 3’ arm, and a TK cassette, and could also be linearized by NotI.

Both targeting vectors were linearized and transfected into the G4 C57/BL6×129 F1 hybrid ES cells (George et al., 2007) using a BTX Electro Square Porator, and positive clones were identified by G418 and FIAU double selection and Southern blot hybridization. Two positive clones for each allele were expanded and injected into C57/BL6 blastocysts to generate chimeric mice which were then mated for germline transmission of the targeted alleles.

### Genotyping

The genotyping of ES cells was carried out by Southern blot hybridization with external probes described before (Fu et al., 2009). Briefly, genomic DNA from G418 and FIAU double-resistant ES clones was digested with BamHI, resolved on 0.8% agarose gels, and transferred onto nitrocellulose membranes. The membranes were then hybridized with 32P labeled external probes. The *Atoh7* detected a 21 kb fragment for the wild-type allele and a 14 kb fragment for the *Atoh7* ^*zsGreenCreERT2*^ allele (Fig. 1A). The *Pou4f2* probe detected a 10 kb fragment for the wild-type allele, and a 6 kb fragment for the *Pou4f2*^*FlagtdTomato*^ allele (see Fig.1B). Once the lines were established, PCR was used for genotyping using primers listed in Supplemental Table 1. PCRs were carried out with the following conditions:: 95 ºC, 1 min, followed by 30 to 35 cycles of 95 ºC, 30 sec; 55 ºC, 30 sec; and 72 ºC, 40 sec, and 5 min of extension chasing at 72 ºC for 5 min.

**Figure 1.**
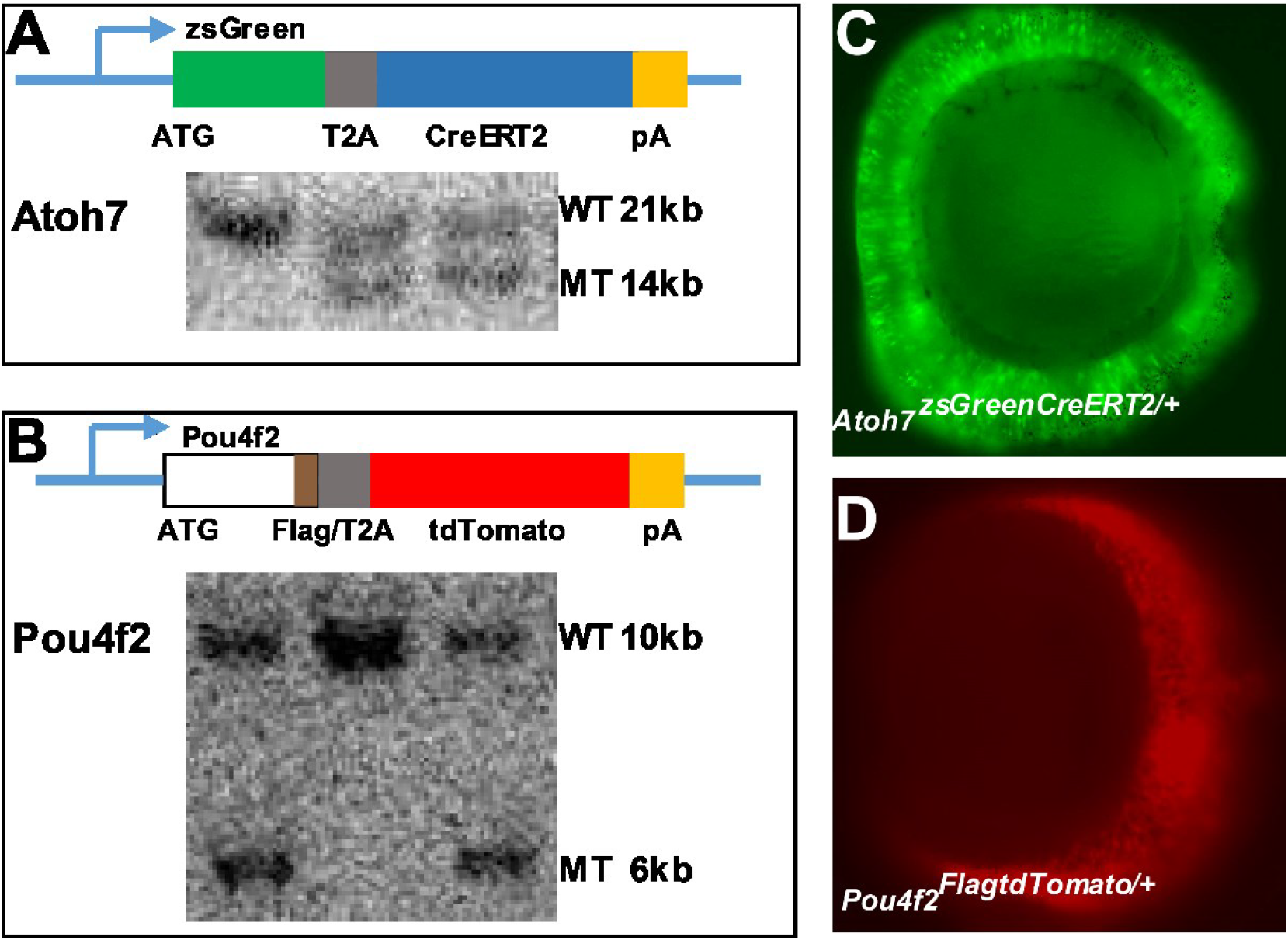
Generation of two knock-in mouse lines, *Atoh7*^*zsGreenCreERT2*^ and *Pou4f2*^*FlagtdTomato*^. **A, B**: Structures of the two lines (diagrams not drawn to scale). In each line, the knock-in cassette starts from the start codon of each native gene. Each cassette expresses two proteins from one ORG separated by the T2A peptide. *Atoh7*^*zsGreenCreERT2*^ expresses zsGreen and CreERT2, whereas *Pou4f2*^*FlagtdTomato*^ expresses a FLAG-tagged version of Pou4f2 and tdTomato. Also shown are genotyping results of ES clones for each allele by Southern hybridization. Clones with two bands carry both the wild-type (WT) and knock-in (MT) alleles. **C. D**: Images of E14.5 *Atoh7*^*zsGreenCreERT2/+*^ (**C**) and *Pou4f2*^*FlagtdTomato/+*^ (**D**) eyes examined by an epifluorescence microscope indicating zsGreen and tdTomaton expression from these two alleles respectively.

### Immunofluorescence Staining

The immunofluorescence staining on mouse embryo or eye sections were performed as previously described (Sapkota et al., 2014; Wu et al., 2015). Briefly, mouse embryos (E12.5-E17.5) or eyes (P0 or older) were collected and fixed with 4% paraformaldehyde for 30 min. After fixation, tissues were rinsed three times with PBST (PBS pH7.4, plus 0.1% Tween 20) for 20 min each time, transferred sequentially into 10%, 20% and 30% sucrose in PBST at 4 ºC until they sank, and then embedded and frozen in Optimum Cutting Temperature (OCT). The frozen tissues were sectioned at 16 µm on a cryostat (Leica CM1850). For immunofluorescence labeling, the section slides were washed for 3 × 10 min with PBST, and blocked with 2% BSA in PBST for 1 hr, then incubated with primary antibodies and fluorescent dye-conjugated secondary antibody and washed with PBST 3 × 10 min. After labeling, the slides were mounted with AquaMount (Thermo Fisher). Antibodies and their dilutions used in this study were: Rabbit anti-Pou4f2 (Santa Cruz, 1:150), Rabbit anti-HA (Santa Cruz, 1:100), Alexa-488 conjugated donkey anti-rabbit IgG (1:400), and Alexa-546 conjugated donkey anti-rabbit IgG (1:400).

### Confocal Imaging and Cell Counting

The images were acquired using a confocal microscope (Leica TCS SP2). When necessary, the contrast of images was adjusted in Adobe Photoshop to the same degree to all images in the same experiment. Cell counting followed a previously reported method (Sapkota et al., 2014). Positive cells for various desired markers (such as zsGreen, tdTomato, HA, Pou4f2) were counted from arbitrary unit lengths in the central regions of retinal sections. At least four sections from different mice were counted.

### Fluorescent flow cytometry analysis of cells

Mouse retinas of the desired developmental stage were dissected out and collected in PBS solution, dissociated by 100 µg/ml trypsin for 10 min at 37 ºC, and then quenched with 100 µg/ml trypsin inhibitor. Cells were harvested by centrifugation at 300 g for 5 min, and re-suspended in PBS for cell sorting on a BD FACS Fusion Cell Sorter. The viable and single cell events were selected for analysis through a forward and side scatter gates. Flow cytometry data were analyzed using the FlowJo (BD Biosciences) analysis software.

### Tamoxifen induced labeling by the Ai9 reporter line

The *Atoh7*^*zsGreenCreERT2*^ mice were bred with the Cre-dependent tdTomato reporter line Ai9 (Madisen et al., 2010) through timed mating. The day when a vaginal plug was detected is set as E0.5. The pregnant mice were intraperitoneally injected of tamoxifen with different doses (0, 4, 8, 80 mg/kg) at E12.5 to fluorescently label progenies of the *Atoh7*-expressing RPCs. Mouse embryos (E14.5) or eyes (P0 or older) were collected for cryo-section and confocal images were obtained as described above. For flat-mount retina imaging, the P0 eyes were fixed by 4% paraformaldehyde in PBS for 1.5 hr and the retinas were dissected from the eyecup carefully, radially incised and flat-mounted on slides with the ganglion cell layer facing up. The slides were mounted with AquaMount for fluorescent microscopy using a Nikon 80i fluorescence microscope equipped with a digital camera and Image Pro analysis software.

## Results

### Generation of the knock-in mouse lines: *Atoh7*^*zsGreenCreERT2*^ and *Pou4f2*^*FlagtdTomato*^

*Atoh7* and *Pou4f2* mark two stages of RGC development. Both the *Atoh7* and *Pou4f2* loci have been targeted multiple times, either by replacing the coding regions with other genes or by adding ectopic tags to the proteins (Badea et al., 2009; Brown et al., 2001; Feng et al., 2010; Fu et al., 2009; Mao et al., 2008; Wang et al., 2000, 2001; Wu et al., 2015; Yang et al., 2003). To study retinal development, we generated two knock-in mouse lines: *Atoh7*^*zsGreenCreERT2*^ and *Pou4f2*^*FlagtdTomato*^ (Fig. 1A and B). In *Atoh7*^*zsGreenCreERT2*^, a cassette coding zsGreen and CreERT2 (Vallier et al., 2001), separated by the viral self-cleaving peptide T2A (Szymczak et al., 2004) was knocked into the *Atoh7* locus and replaced the *Atoh7* ORF by homologous recombination. For the *Pou4f2*^*FlagtdTomato*^ allele, a cassette expressing the FLAG tag followed by a T2A sequence and sequences encoding tdTomato was knocked into the *Pou4f2* locus in frame right after the last codon. Based on prior experiences, such modifications of these two alleles should not interfere with the promoter activities of the two genes (Badea et al., 2009; Fu et al., 2009; Wu et al., 2015). Thus, *Atoh7*^*zsGreenCreERT2*^ was a null allele but designed to express both zsGreen and CreERT2 from the *Atoh7* promoter (Fig. 1A), whereas *Pou4f2*^*FlagdTomato*^ was supposed to be a functional wild-type allele expressing a FLAG-tagged version of Pou4f2 and tdTomato from the *Pou4f2* promoter (Fig. 1B). The reason to create a null *Atoh7* allele was that it will allow us to study the *Atoh7*-expressing RPCs in both wild-type (heterozygous) and mutant background, which would be essential for probing the mechanism by which Atoh7 function in RGC development. On the other hand, we decided to create a functional fluorescently labeled *Pou4f2* allele because most *Pou4f2*-null RGCs die by apoptosis (Gan et al., 1999; Xiang, 1998), making it difficult to study the mutant cells. Addition of the FLAG tag to Pou4f2, although not tested yet in this study, also generated another tool for studying the function of Pou4f2. Both targeting constructs were transfected into the G4 mouse ES cells (George et al., 2007) by electroporation, and the ES cell clones were identified by Southern hybridization using external probes for each allele (Fu et al., 2009). Cells from two correctly targeted *Atoh7*^*zsGreenCreERT2*^ and *Pou4f2*^*FlagtdTomato*^ ES clones each were injected into BL/6C57 blastocysts and the targeted alleles were successfully transmitted through the germline to establish both mouse lines. Although a *Neo* cassette flanked by two *loxP* sites was present in both alleles (not shown in the diagrams in Fig. 1A and B), it was not deleted in this study since it appeared not to interfere with the correct expression of either allele (Fu et al., 2009). Homozygous *Atoh7*^*zsGreenCreERT2/zsGreenCreERT2*^ and *Pou4f2*^*FlagtdTomato/FlagtdTomato*^ mice appeared largely normal. Nevertheless, eyes of *Atoh7*^*zsGreenCreERT2/zsGreenCreERT2*^ mice lacked the optic nerve, whereas those of *Pou4f2*^*FlagtdTomato/FlagtdTomato*^ mice had normal optic nerves (data not shown). These results indicated that, as designed, *Atoh7*^*zsGreenCreERT2*^ was a null allele, and *Pou4f2*^*FlagtdTomato*^ was a functional wild-type allele. Under an epifluorescence microscope, E14.5 *Atoh7*^*zsGreenCreERT2/+*^ and *Pou4f2*^*FlagtdTomato/+*^ embryos emitted green and red fluorescence respectively from the retina, indicating the knock-in cassettes were expressed as designed (Fig. 1C, D).

### zsGreen expression in the *Atoh7*^*zsGreenCreERT2/+*^ retina

The purpose of generating these two alleles was to label the cells expressing the two transcription factors. It was thus critical for the fluorescent proteins to faithfully represent the expression of the native proteins. Hence, we examined zsGreen expression pattern in the *Atoh7*^*zsGreenCreERT2/+*^ mouse by directly detecting its fluorescence on retinal sections from different developmental stages by confocal microscopy. During development, Atoh7 is expressed in a subset of RPCs from E11 to P0 with a peak at around E14.5 (Brown et al., 1998; Fu et al., 2009; Miesfeld et al., 2018). As expected, we observed similar dynamic expression patterns of zsGreen in the *Atoh7*^*zsGreenCreERT2/+*^ retina throughout development (Fig. 2A-D). At E12.5, zsGreen was expressed in a subset of RPCs in the central retinal region (Fig. 2A), a pattern essentially identical to the native Atoh7. At E14.5, zsGreen expression reached the peak and had spread across the whole retina (Fig. 2B, E, E’). At this stage, whereas zsGreen-expressing cells were mostly found in the neuroblast layer (NBL), which were RPCs, some were also observed in the inner ganglion cell layer (GCL). This was different from the native Atoh7 whose expression is confined to the NBL. This difference was likely due to the stability of the zsGreen protein. Expression of zsGreen in RPCs in the NBL decreased markedly at E17.5 (Fig. 2C), and only very few zsGreen-expressing RPCs were observed at P0 (Fig. 2D), but no zsGreen-expressing RPCs were detected at P5 (data not shown). On the other hand, at all these stages, particularly at E17.5 and P0, zsGreen expression could be observed in differentiated cells including RGCs in the GCL and photoreceptors in the outmost side (future out nuclear layer) where expression of the native Atoh7 is inactivated (Fig. 2C and D). Since *Atoh7*-expressing cells adopt multiple retinal cell fates (Brzezinski et al., 2012; Feng et al., 2010; Yang et al., 2003), the differentiated zsGreen-expressing cells were likely progenies of Atoh7-expressing RPCs. Retinal sections from E14.5 *Atoh7*^*zsGreenCreERT2/lacZ*^ embryos also expressed zsGreen extensively (Fig. 2H), but no obvious GCL was observed. Since *Atoh7*^*lacZ*^ is a null allele (Wang et al., 2001), this observation further confirmed that *Atoh7*^*zsGreenCreERT2*^ was also a true null allele.

**Figure 2.**
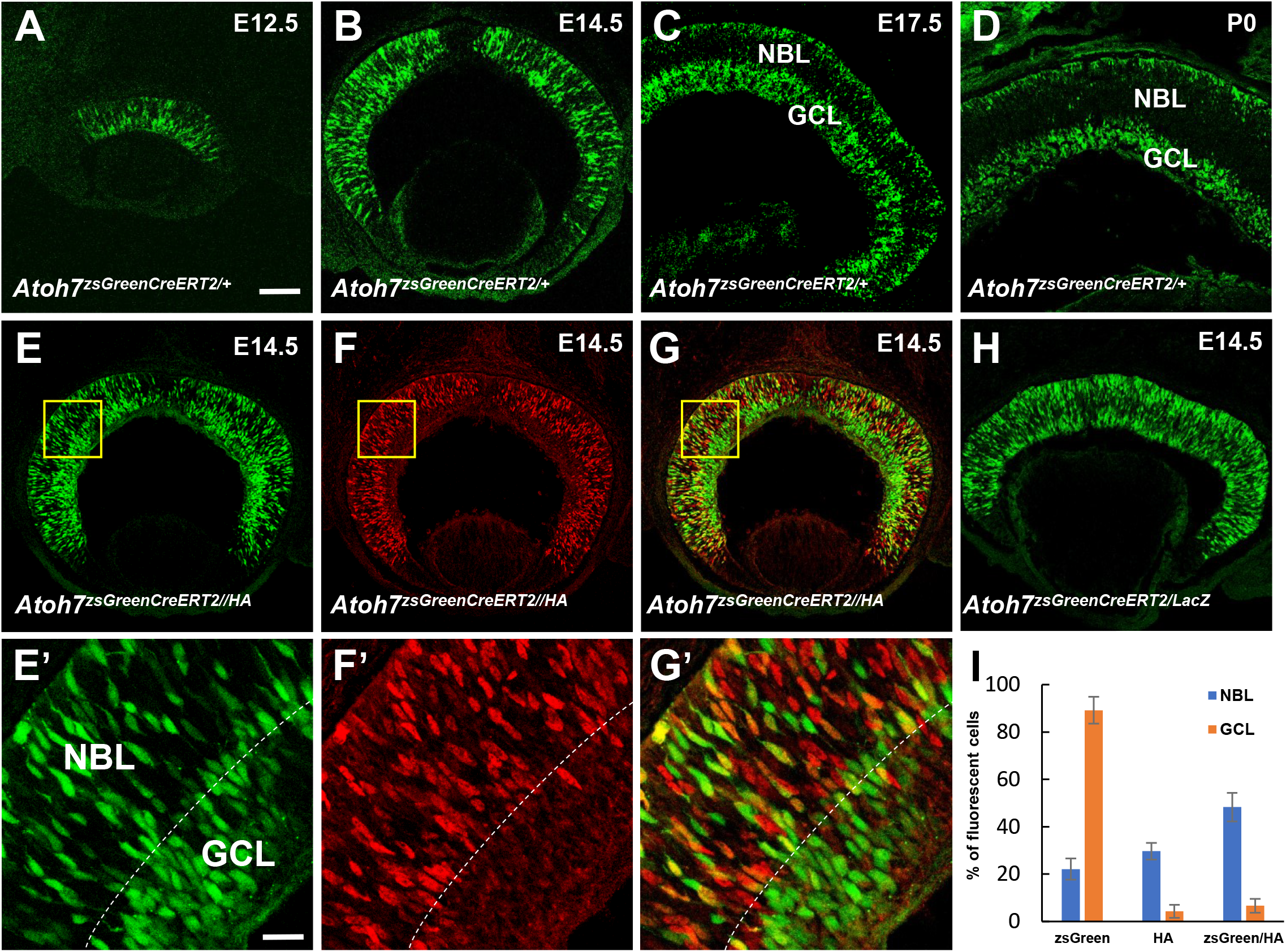
zsGreen expression in the *Atoh7*^*zsGreenCreERT2/+*^ retina. **A-D:** Confocal images displaying zsGreen (green) expression in the *Atoh7*^*zsGreenCreERT2/+*^ retinal sections from E12.5 (**A**), E14.5 (**B**), E17.5 (**C**) and P0 (**D**) embryos. **E-G and E’-G’:** Comparison of zsGreen expression (green) with Atoh7^HA^ (red) in the E14.5 *Atoh7*^*zsGreenCreERT2/HA*^ retina by immunofluorescence staining with an anti-HA antibody. **E’-G’** are high-magnification images of the boxed areas in **E-G**. The boundary between the neuroblast layer (NBL) and the ganglion cell layer (GCL) is indicated by a dotted line. **H:** Retinal section from E14.5 *Atoh7*^*zsGreenCreERT2/lacZ*^ embryo which is null for *Atoh7* displays expression of zsGreen. **I:** Quantification of zsGreen and Atoh7^HA^ positive cells in the NBL and GCL. Error bars indicate ± SD. Scale bars: in **A** for A–H, 200 µm; in E’ for E’-G’,40 µm.

To further examine how the *Atoh7*^*zsGreenCreERT2/+*^ allele represented the wild-type allele, we compared zsGreen expression with Atoh7^HA^ in the E14.5 *Atoh7*^*zsGreenCreERT2/HA*^ retina by immunofluorescence staining with an anti-HA antibody. The *Atoh7*^*HA*^ allele tags the Atoh7 protein with a hemagglutinin epitope and recapitulates the wild-type protein expression completely (Fu et al., 2009; Miesfeld et al., 2018). As mentioned above, whereas Atoh7^HA^ was detected mostly in the NBL where RPCs resided, zsGreen was found in both the NBL and GCL (Fig. 2E-G, E’-G’). In the NBL, zsGreen and Atoh7^HA^ overlapped significantly; of all the cells expressing either zsGreen or Atoh7^HA^, 48% expressed both, 22% expressed just zsGreen, and 30% expressed just Atoh7^HA^ (Fig. 2I). The differences of expression from these two alleles were likely due to the intrinsic properties of the proteins, instead of the differential activities of the promoters in the two alleles. The Atoh7^HA^ only cells in the NBL were likely those just starting to express Atoh7, but the zsGreen protein had not matured yet to emit fluorescence. The zsGreen only cells were likely those that had turned off *Atoh7*, but zsGreen persisted in them due to its stability. Because of this stability of zsGreen, in the GCL where the *Atoh7* is turned off, very few cells expressed Atoh7^HA^, but many cells still expressed zsGreen (Fig. 2E-G, E’-G’, I). These results indicated that the *Atoh7*^*zsGreenCreERT2*^ allele largely recapitulated the expression of Atoh7 in RPCs, but labeled cells of a slightly delayed time window; thus zsGreen positive cells also included recent progenies from the Atoh7-expressing RPCs.

### tdTomato from the *Pou4f2*^*FlagtdTomato*^ allele labels RGCs

The *Pou4f2*^*FlagtdTomato*^ allele was generated to label RGCs, since *Pou4f2* is one of the earliest RGC marker genes (Xiang et al., 1995). To assess whether tdTomato from *Pou4f2*^*FlagtdTomato*^ indeed marked RGCs, we examined its expression pattern on *Pou4f2*^*FlagtdTomato/+*^ retinal sections from different developmental stages. The tdTomato fluorescence was detected also directly by confocal imaging on retinal sections. Pou4f2 starts to be expressed at E11.5, and continue to be expressed in all later stages, although it is expressed in only a subset of RGCs in postnatal retinas (Badea et al., 2009; Xiang et al., 1995). Consistent with these patterns, we detected tdTomato in the GCL on *Pou4f2*^*FlagtdTomato*^ retinas from E12.5 to P5 (Fig. 3A-E). The temporal dynamics of tdTomato mirrored that of Pou4f2 very closely. At E12.5, tdTomato expression was confined to the GCL of the central retinal region (Fig. 3A). From E14.5 to P0, it continued to be expressed in most, if not all, cells, in the GCL at high levels (Fig. 3B and C, and data of P0 not shown). By P5, however, only a fraction of RGCs expressed tdTomato at high levels (Fig. 3D). It should be noted that unlike Pou4f2, tdTomato was not confined to the nucleus; therefore the axons in the neural fiber layer were also labeled (Fig. 3B and C). This was further illustrated by flat-mount imaging of the whole retina at P16, which displayed not just the RGC somas, but also the radially projecting axons converging at the optic disc (Fig. 3E).

**Figure 3.**
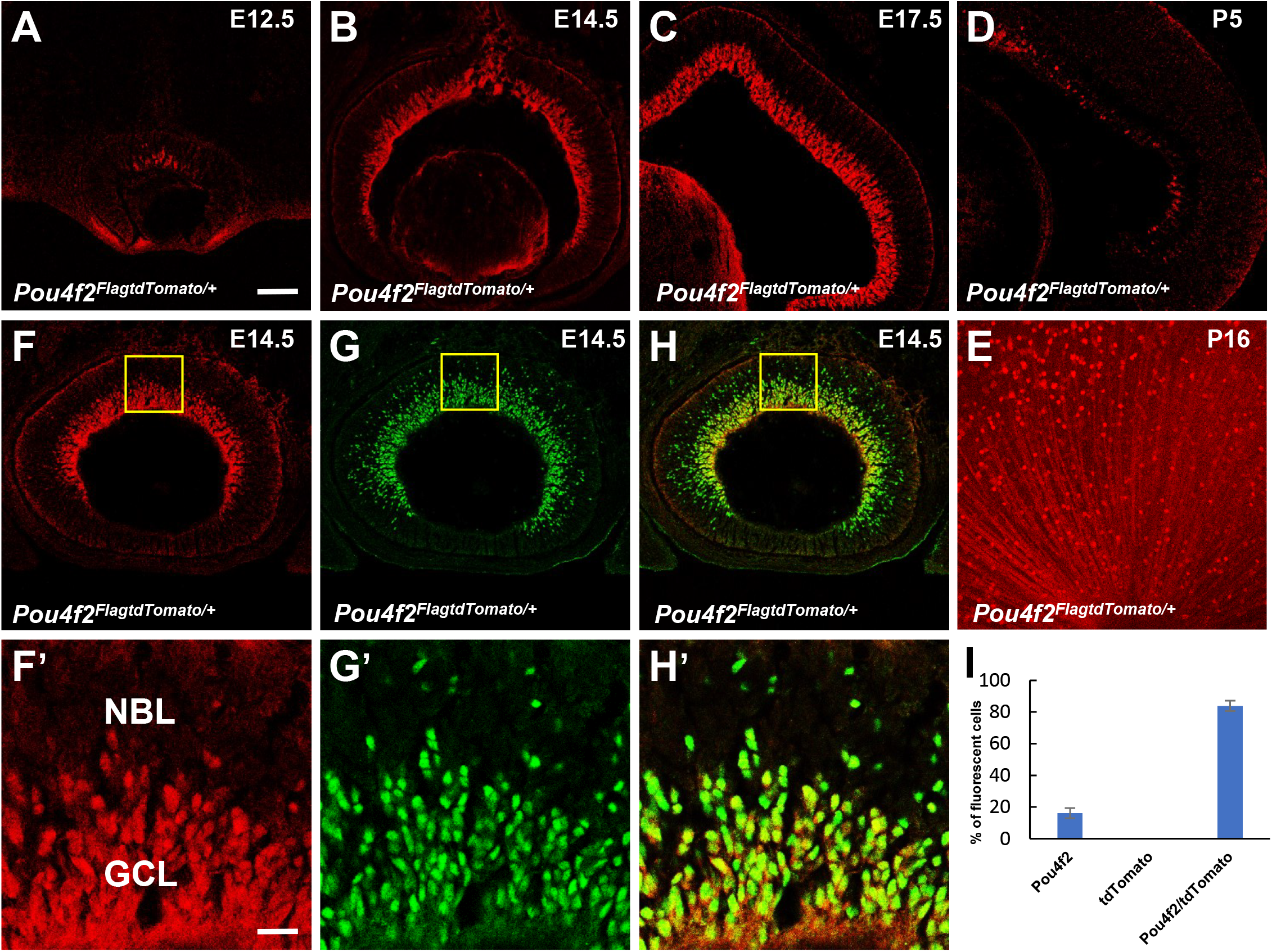
tdTomato from the *Pou4f2*^*FlagtdTomato*^ allele labels RGCs. **A-D:** Retinal sections of *Pou4f2*^*FlagtdTomato/+*^ from E12.5 (**A**), E14.5 (**B**), E17.5 (**C**) embryos P5 (**D**) pups displayed tdTomato (red) expression in the ganglion cell layer**. E.** Flatmount image of P16 *Pou4f2*^*FlagtdTomato/+*^ retina showing tdTomato expression in RGC somas and axons. **F-I, F’-H’:** Comparison of tdTomato (red) expression with Pou4f2 (green) in the E14.5 *Pou4f2*^*FlagtdTomato/+*^ retina by immunofluorescence staining with an anti-Pou4f2 antibody. **F’-H’** are high-magnification images of the boxed areas in **F-H**. **I:** Quantification of tdTomato and Pou4f2 positive cells in the E14.5 *Pou4f2*^*FlagtdTomato/+*^ retina. Error bars indicate ± SD. Scale bars: in **A** for **A–E**, 200 µm; in **F’** for **F’-H’**,40 µm.

In order to further evaluate how the *Pou4f*^*FlagtdTomato*^ allele faithfully represented the native *Pou4f2* allele, we performed immunofluorescence co-staining with an anti-Pou4f2 antibody on *Pou4f2*^*FlagtdTomato/+*^ retinal sections at E14.5, which showed that Pou4f2 and tdTomato highly overlapped (Fig. 3F-H, F’-H’). tdTomato and Pou4f2 had almost identical expression patterns in the GCL. Counting of tdTomaton- and Pou4f2-expressing cells revealed that more than 80% of positive cells expressed both proteins (Fig. 3I). Interestingly, about 20% of Pou4f2 positive cells did not show tdTomato fluorescence and almost all these cells were located in the NBL (Fig. 3G, H). These cells were likely newly born RGCs that had not migrated to the GCL. Thus, similar to the zsGreen protein from the *Atoh7*^*zsGreenCreERT2*^ allele, the delayed expression of tdTomato likely reflected the time lapse from translation of the protein to maturation required for fluorescence emission, rather than inherent differences in the activities of the two alleles. Consistent with this idea, essentially all tdTomato-expressing cells expressed Pou4f2. Thus, the normal regulatory mechanism for *Pou4f2* expression was maintained in *Pou4f2*^*FlagtdTomato*^ allele and the red fluorescence from tdTomato is a good reporter of *Pou4f2* promoter activity in RGCs, although with some time delay.

### *Atoh7*^*zsGreenCreERT2*^ and *Pou4f2*^*flagtdTomato*^ mark two consecutive phases of the RGC developmental trajectory

Atoh7 marks a subset of RPCs competent for the RGC lineage, whereas Pou4f2 expression coincides with the fate commitment of RGCs. These two factors overlap transiently in cells that have just turned on *Pou4f2* and have not turned off *Atoh7* (Fu et al., 2009). Thus, Atoh7 and Pou4f2 mark two consecutive phases along the developmental trajectory of the RGC lineage. To further examine how *Atoh7*^*zsGreenCreERT2*^ and *Pou4f2*^*FlagtdTomato*^ represented these two phases, we examined the expression of zsGreen and tdTomaton in the E14.5 *Atoh7*^*zsGreenCreERT2/+*^;*Pou4f2*^*FlagtdTomato/+*^ retina. Similar to the native Atoh7 and Pou4f2 proteins, zsGreen and tdTomato were detected in two largely separate domains, zsGreen largely in the NBL whereas tdTomation mostly in the GCL, but with significant overlap (Fig. 4). However, unlike the native Atoh7 and Pou4f2 whose overlap largely occurs in the NBL and marks newly generated RGCs that have not migrated to the GCL (Fu et al., 2009), cells expressing both zsGreen and tdTomato were observed mostly in the GCL. In the GCL, two populations of cells were observed; some expressed both zsGreen and tdTomato whereas others expressed just tdTomato (Fig. 4C, C’). The difference between the native proteins and the fluorescent proteins was likely caused by the maturation lapse and stability of the zsGreen and tdTomato proteins as discussed above, leading to a delayed time window for these proteins to be detected. Thus, cells expressing both zsGreen and tdTomato in the GCL were likely newly arrived RGCs from the NBL, whereas cells expressing tdTomato only in the GCL were generated earlier. Nevertheless, *Atoh7*^*zsGreenCreERT2*^ and *Pou4f2*^*FlagtdTomato*^ still allowed us to label the two critical phases of RGC development. As discussed later, the early stages when zsGreen and tdTomato could not be efficiently detected by confocal imaging here may still be captured by lowering the detection threshold in other techniques such FACS to study these two phases of RGC development.

**Figure 4.**
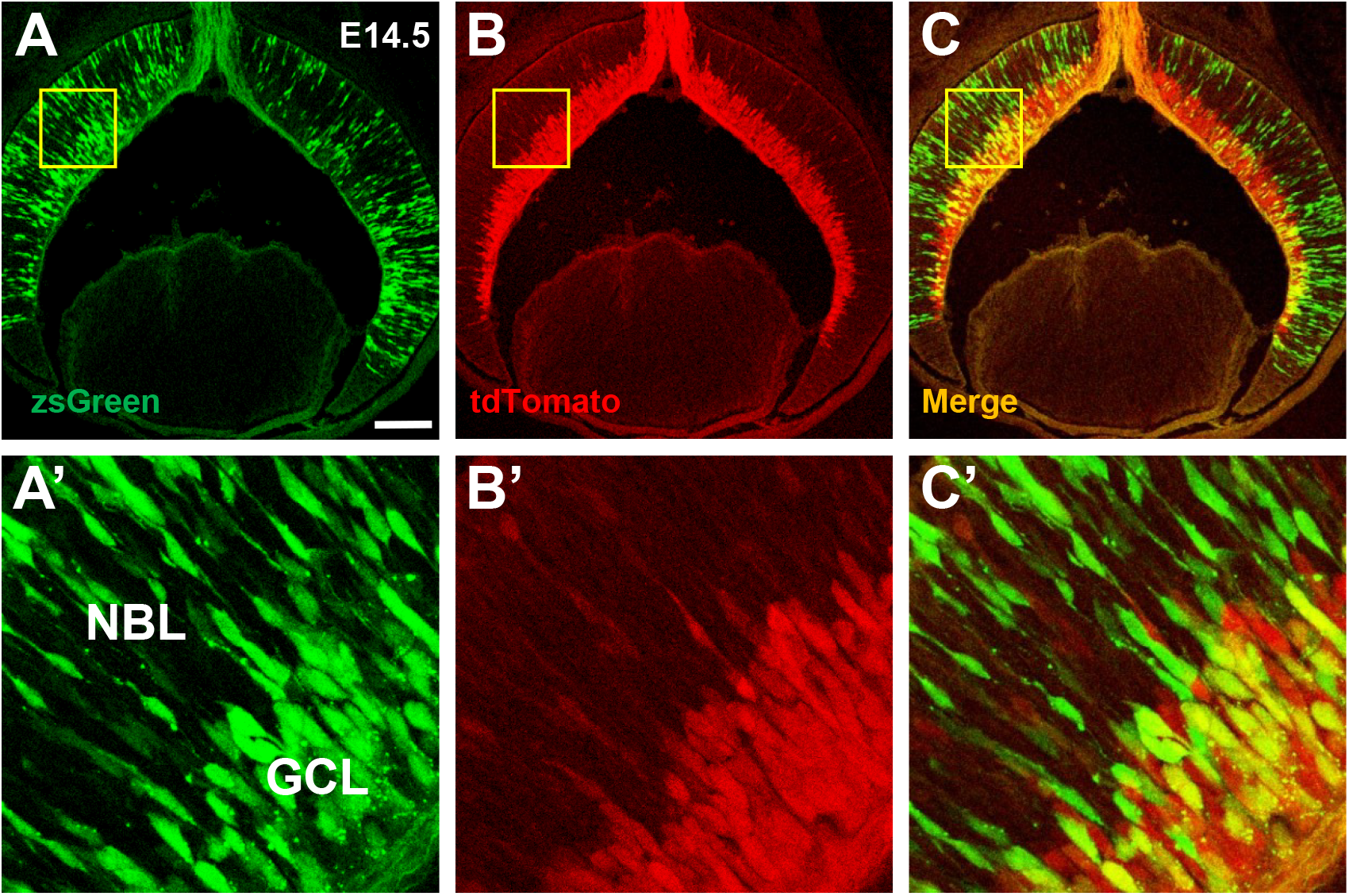
*Atoh7*^*zsGreenCreERT2*^ and *Pou4f2*^*FlagtdTomato*^ mark two consecutive phases of RGC development. **A-C:** Retinal section from E14.5 *Atoh7*^*zsGreenCreERT2/+*^;*Pou4f2*^*FlagtdTomato/+*^ embryo displaying the expression of the zsGreen (green) and tdTomato (red). **A’-C’:** High-magnification images of the boxed areas in **A-C**. The NBL and GCL layers are indicated in image **A’**. Note that the overlap of zsGreen and tdTomato occurs in the GCL. Scale bars: in **A** for **A–C**, 200 µm; in **A’** for **A’-C’**, 40 µm.

### Isolation of Atoh7- and Pou4f2-expressing cells by FACS

A prerequisite for studying the properties of different cell states/types during development is the ability to isolate the cell populations to purity. The *Atoh7*^*zsGreenCreERT2*^ and *Pou4f2*^*FlagtdTomato*^ could potentially be used to isolate cells of the two critical phases of RGC development by fluorescence-activated cell sorting (FACS). To assess this possibility, we treated *Atoh7*^*zsGreenCreERT2/+*^;*Pou4f2*^*FlagtdTomato/+*^ retinas with trypsin and were able to dissociate them into single cell suspensions. As observed on retinal sections, cells expressing zsGreen or tdTomato alone, or both, could be readily observed in the suspensions (Fig. 5A-C). We then subjected the dissociated E14.5 *Atoh7*^*zsGreenCreERT2/+*^ and *Pou4f2*^*FlagtdTomato/+*^ retinal cells to FACS. To be able to isolate cells that had just begun to express the two fluorescent proteins, we set the gating threshold for the fluorescence to relatively low levels. As shown in Fig. 5D and E, zsGreen- and tdTomato-positive cells were well separated from the negative cells by FACS. In the experiment shown, we were able to isolate 28.3% zsGreen-positive and 13.9% tdTomato-positive cells from E14.5 *Atoh7*^*zsGreenCreERT2/+*^ and *Pou4f2*^*FlagtdTomato/+*^ retinas respectively. Similar proportions of cells were obtained in multiple repeated experiments with E14.5 retinas, and zsGreen and tdTomaton positive cells could also be sorted out from other stages (e.g. E12.5 and E17.5) (data not shown). The purified cells were highly viable; more than 90% of them were alive as determined by trypan blue staining (data not shown). Since these cells were enriched for specific phases of RGC development, they could be valuable resources for further studying RGC development using such techniques as single cell RNA-seq (Clark et al., 2019; Shekhar et al., 2016) and single cell ATAC-seq (Satpathy et al., 2019).

**Figure 5.**
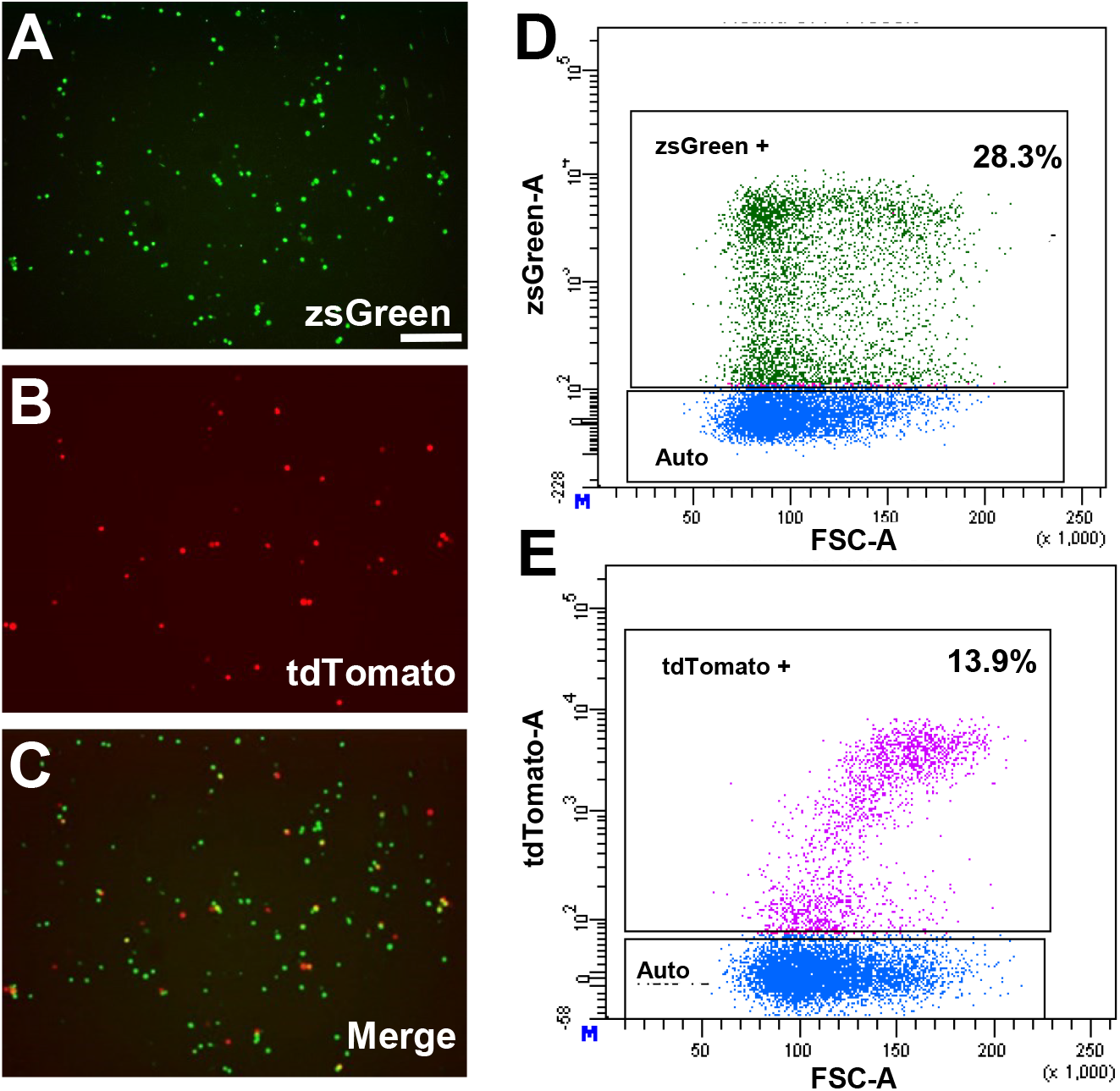
Isolation of *Atoh7*- and *Pou4f2*-expressing cells by FACS. **A-C:** Single cell suspensions dissociated from E14.5 *Atoh7*^*zsGreenCreERT2/+*^;*Pou4f2*^*FlagtdTomato/+*^ retinas express zsGreen or tdTomato alone, or both (see yellow cells in **C**). **D, E:** zsGreen (**D**) and tdTomato (**E**) positive cells from *Atoh7*^*zsGreenCreERT2/+*^ and *Pou4f2*^*FlagtdTomato/+*^ retinas were sorted by FACS respectively. The gating threshold for the fluorescence was set to relatively low levels in order to isolate cells that had just begun to express the two fluorescent proteins. Y axis is fluorescence intensity of the fluorescent proteins, and X axis is autofluorescence.

### Permanently labeling progenies of *Atoh7* positive RPCs using the Ai9 reporter

The *Atoh7*^*zsGreenCreERT2*^ allele was designed for dual purposes, to both mark *Atoh7*-expressing cells directly and to serve as a tamoxifen-inducible deleter in cells expressing Atoh7, since the knock-in cassette encodes both zsGreen and CreERT2 in the same ORF linked by T2A. CreERT2 is a fusion protein of Cre recombinase and a mutant version of the estrogen receptor hormone binding domain, which is retained in the cytoplasm and inactive, but translocates to the nucleus and becomes active upon tamoxifen binding (Vallier et al., 2001). Thus, if CreERT2 was expressed and functioned as designed, it could be used in various ways to study gene functions and different aspects of retinal development. One application of this allele was to trace the fate of Atoh7-expressing RPCs. To test the function of this allele, we crossed it with the Ai9 reporter line, which expresses tdTomato constitutively following Cre-mediated recombination (Madisen et al., 2010). Pregnant dams carrying *Atoh7*^*zsGreenCreERT2/+*^;*Ai9* were treated at E12.5 with a series of doses of tamoxifen (0, 4, 8, and 80 mg/kg). Embryos (E14.5) or pups (P0 or older) from these dams were then collected and analyzed for tdTomato fluorescence (Fig. 6). At both E14.5 and P0, a dose-dependent response of tdTomato-expressing cells to tamoxifen was observed (Fig. 6A-D, G-J). In the control retina (0 mg/kg), only very scarce tdTomato-positive cells could be seen, indicating that there was only a very low level of leaky activity of CreERT2 (Fig. 6A and G). As the tamoxifen dosage increased, there were increased numbers of tdTomato-positive cells which mostly located in the GCL (Fig. 6B-D). In E14.5 retinas treated with the highest dosage (80 mg/kg), most cells in the GCL expressed tdTomato, and they were largely zsGreen-positive, indicating that they were indeed progenies of Atoh7-expressing RPCs (Fig. 6D-F). Expression of tdTomato persisted in postnatal (P16 and P30) retinas with no noticeable reduction in the intensity of fluorescence or the number of positive cells (Fig. 6K and data not shown). Thus, these cells were indeed permanently labeled by tdTomato and the ectopic fluorescent protein was not toxic to the cells. In addition, tdTomato cells were not only found in the GCL, but also in other layers of both the E14.5 and P16 retinas (Fig. 6D and L). Based on their locations and morphologies, tdTomato-positive cells in the other layers included photoreceptors, horizontal cells, and amacrine cells (Fig. 6L). These observations were consistent with previous reports that *Atoh7*-expressing RPCs adopt multiple retinal cell fates.

**Figure 6.**
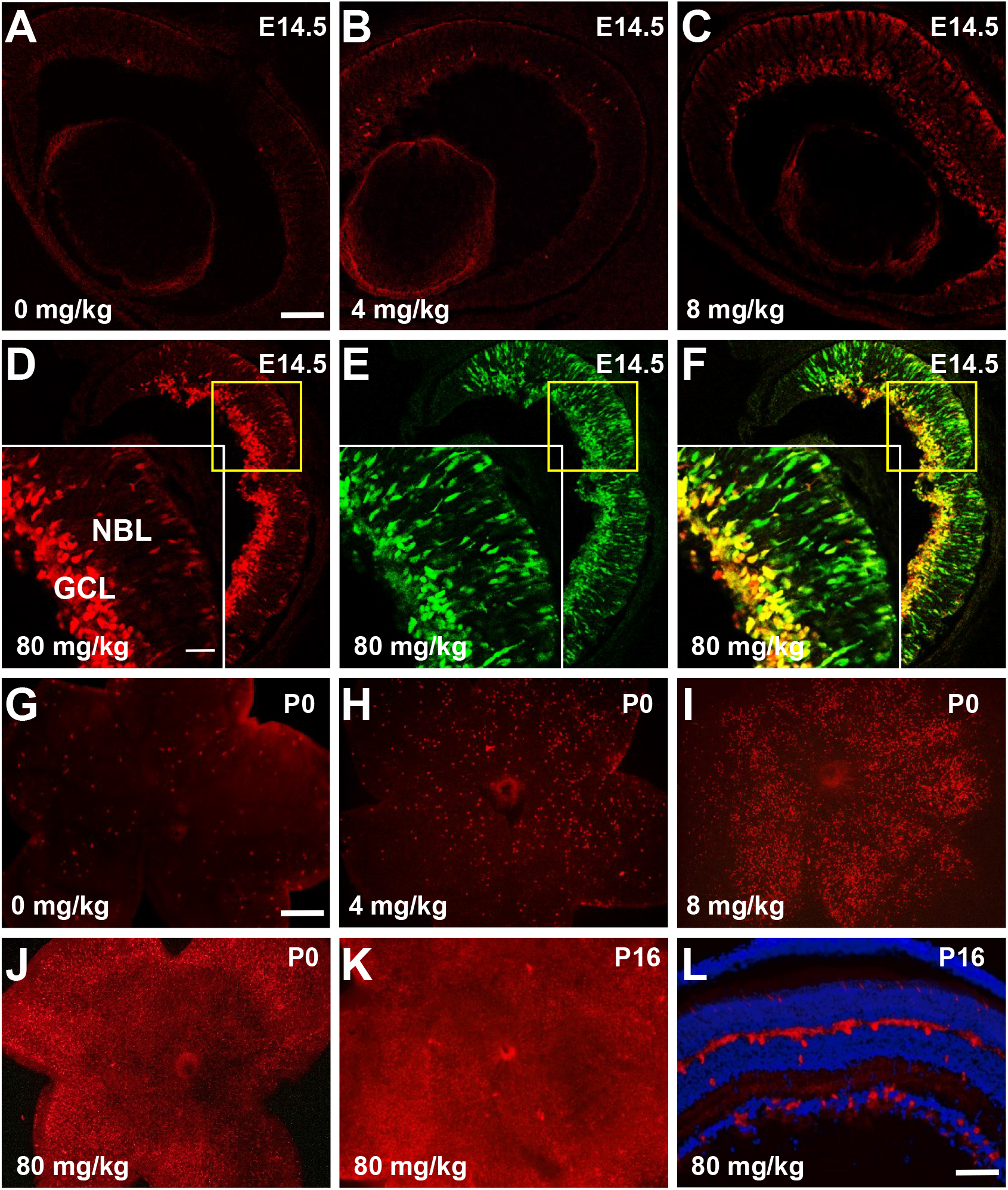
Permanently labeling *Atoh7* positive RPC progenies by the Ai9 reporter. **A-D:** tdTomato positive cells from E14.5 *Atoh7*^*zsGreenCreERT2/+*^;*Ai9* retinas after administration of tamoxifen at E12.5. There is increased tdTomato positive cells as tamoxifen dosage increases. **E-F**: Comparison of zsGreen to tdTomato in the E14.5 *Atoh7*^*zsGreenCreERT2/+*^;*Ai9* retina after treatment with tamoxifen at high dosage (80 mg/kg). Inlets are high-magnification images of the yellow boxed areas. Note tdTomato is delayed as compared to zsGreen and their overlaps largely occur in the GCL. (**G-J**) tdTomato positive cells from P0 *Atoh7*^*zsGreenCreERT2/+*^;*Ai9* retinas after administration of tamoxifen at E12.5. Similar dosage response can be seen as observed at E14.5. **K:** tdTomato positive cells in P16 *Atoh7*^*zsGreenCreERT2/+*^;*Ai9* retina indicating the fluorescent cell persist in later stages. **L.** P16 *Atoh7*^*zsGreenCreERT2/+*^;*Ai9* retinal section indicating progenies (red) of Aoh7-cells include all early born retinal cell types: cones, horizontal cells, amacrine cells, and RGCs. Blue is DAPI staining of the nuclei. Scale bars: in A for A–F, 200 µm; in D inlet for inlets in D-F, 70 µm; in G for G-K, 500 µm; in L, 100 µm.

## Discussion

In this report, we describe two novel knock-in alleles, *Atoh7*^*zsGreenCreERT2*^ and *Pou4f2*^*FlagtdTomato*^, that could be very useful tools for studying the development and biology of RGCs. These two alleles make use of the fact *Atoh7* and *Pou4f2* are expressed in two consecutive phases of RGC development. We show that the knock-in cassettes encoding fluorescent proteins are expressed in patterns highly similar to those of the respective native genes, although with some delay due to the maturation time for the fluorescent proteins. These two alleles thus add to an extensive list of genetically modified alleles of these two loci. The strong fluorescence of zsGreen and tdTomato from these two alleles and the dual-purpose design of *Atoh7*^*zsGreenCreERT2*^ make them very useful in many potential applications.

First, these alleles enable the specific and efficient isolation of *Atoh7*-expressing and *Pou4f2*-expressing cells from different retinal developmental stages. These cells will allow for further investigation of their properties at both transcriptomic and epigenetic levels. Since *Atoh7*-expressing cells can adopt different cell fates, they likely are not a homogeneous population. However, *Atoh7* is only expressed in a small subset of RPCs, making it hard to characterize them. Being able to isolate the *Atoh7*-expressing cells should significantly facilitate the study and provide insight into their heterogeneity and the epigenetic underpinning for their ability to adopt different development trajectories. Since most *Atoh7*-expressing RPCs are at the last cell cycle, the indirect labeling method used previously did not capture the stage when *Atoh7*-expressing cells are still dividing and their fate not determined (Gao et al., 2014). Since *Atoh7*^*zsGreenCreERT2*^ expresses zsGreen from its promoter directly and zsGreen overlaps with the native protein significantly, we should be able to capture most of the time window of *Atoh7* expression. The lapse caused by zsGreen maturation may be mitigated by lowering the gating threshold when isolating the cells by FACS. In addition, high stability of zsGreen may be an advantage, since it will also enable the isolation of recently born progenies of *Atoh7*-expressing RPCs, providing continuity in the different cell lineages. Similarly, being able to isolate *Pou4f2*-expressing cells with the *Pou4f2*^*FlagtdTomato*^ allele will make it possible to study RGCs after their fate is committed. One critical question that can be addressed using these cells is the specific transcription program activated during RGC fate specification and differentiation. Since *Atoh7*^*zsGreenCreERT2*^ and *Pou4f2*^*FlagtdTomato*^ label two separate but overlapping populations of cells along the RGC developmental trajectory, transcriptomic and epigenomic studies of these cells using single cell technology should allow us to decipher changes in the epigenetic landscape required for the transition from multipotent RPCs to fate committed RGCs. Further, since *Pou4f2*^*FlagtdTomato*^ labels RGCs throughout development and persists postnatally, it is also possible to isolate RGCs from different stages and use them to study the genetic programs involved in RGC maturation and subtype specification, which are significant but currently poorly studied issues. Additionally, since the two proteins can be observed in live cells, it may also be possible to monitor RGC formation directly using retinal organoid cultures derived from embryonic stem cells carrying these alleles (Eiraku et al., 2011). Such observations may lead to insights into cell division, cell migration, and morphological changes of RGCs during development and alterations in these aspects caused by various experimental perturbations. These alleles may also be used to directly monitor RGCs in mouse models of retinal diseases such as glaucoma in which RGCs are implicated.

Our results indicate that the *Atoh7*^*zsGreenCreERT2*^ allele also functions as an efficient and tamoxifen dose-dependent deleter for various Cre-LoxP mediated genetic recombination applications. In combination with a conditional reporter (e.g. Ai9), this can be used as an alternative means to label RGCs and other progenies of *Atoh7*-expressing cells. The allele may also be used as a deleter for conditional knockout of floxed gene alleles in a time-specific and dose-dependent fashion to study their functions in the retina, particularly in RGCs. Also, although previous studies using regular Cre have revealed that *Atoh7*-expressing cells adopt multiple retinal cell fates (Brzezinski et al., 2012; Feng et al., 2010; Yang et al., 2003), the behavior of individual RPCs, such as the number and cell type composition of progenies they produce, have not been definitely addressed. The dose-dependent nature of CreERT2 will enable us to unequivocally address this question by sparsely labeling individual Atoh7 clones at low doses of tamoxifen.

The *Atoh7*^*zsGreenCreERT2*^ and *Pou4f2*^*FlagtdTomato*^ alleles may also serve as useful tools to monitor effort in generating RGCs, both in vitro and in vivo, for developing cell replacement therapies. RGCs are the major cell type affected in glaucoma and a number of other retinal diseases. Since normally RGCs do not regenerate, damages to RGCs are usually permanent. As in other fields of stem cell biology, much effort is being invested in developing cell replacement measures to treat these diseases. RGCs have been generated in vivo from embryonic stem cells, but often with low efficiency (Gill et al., 2014). Using mouse stem cells carrying both alleles or just *Pou4f2*^*FlagtdTomato*^, it may be possible to directly monitor the generated of RGCs without other labeling techniques, which may streamline the process of developing new procedures and improving efficiencies.

## Acknowledgments

We thank other members of the Mu laboratory and members of the Department of Ophthalmology and the Developmental Genomics Group, University of Buffalo, for helpful discussions. The knock-in mouse lines were created by the Gene Targeting and Transgenic Resource Core at the Roswell Park Cancer Institute. Cell sorting was performed at the Department of Flow & Image Cytometry at the Roswell Park Comprehensive Cancer Center. This project was supported by grants from the BrightFocus Foundation (G2016024) and the National Eye Institute (EY020545) to X.M.

---

**Supplementary Table 1.**
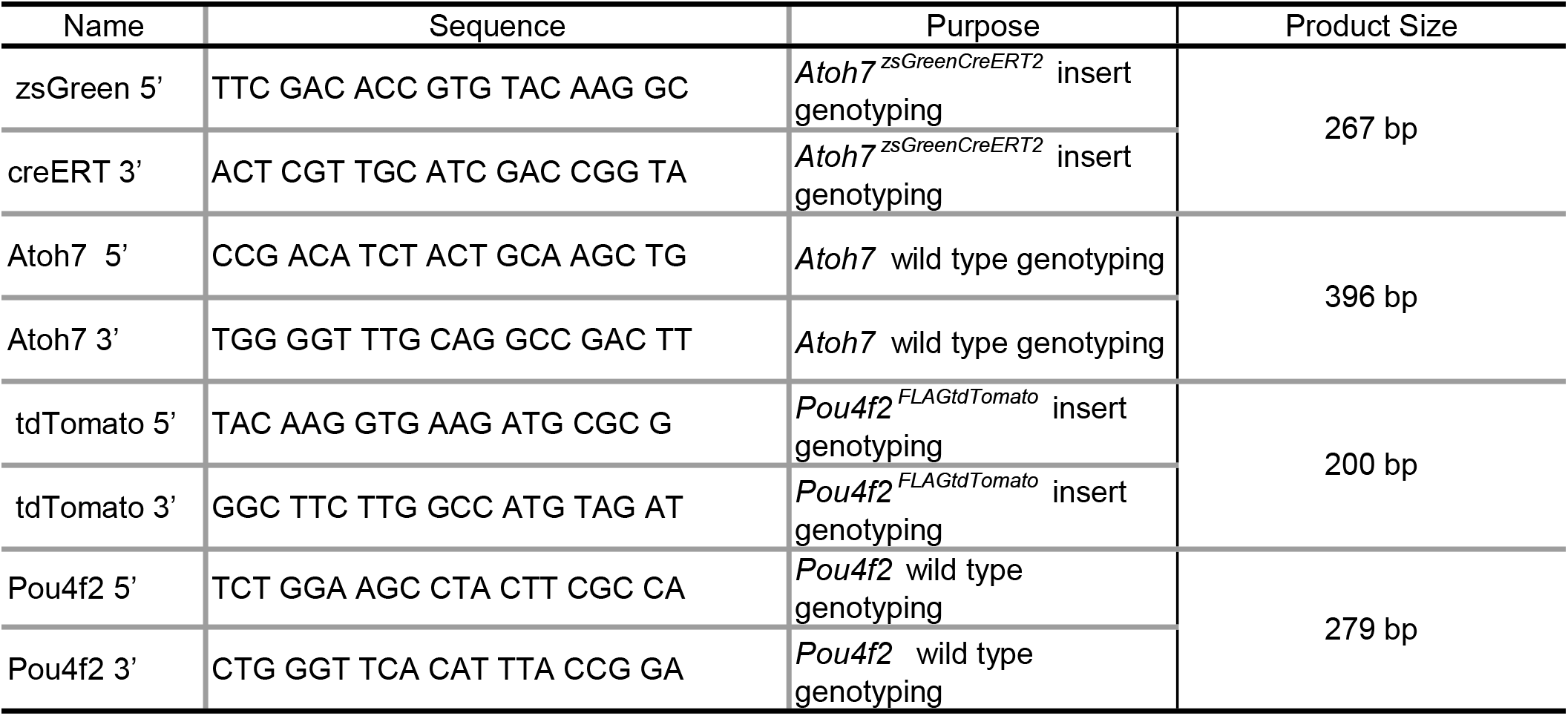
List of genotyping primers

